# CertPrime: a new oligonucleotide design tool for gene synthesis

**DOI:** 10.1101/2025.04.29.650686

**Authors:** David Luna-Cerralbo, Ana Serrano, Irene Blasco-Machín, Fadi Hamdan, Juan Martínez-Oliván, Esther Broset, Pierpaolo Bruscolini

**Author notes:** Corresponding authors (Esther Broset), (Pierpaolo Bruscolini). Email addresses:* (David Luna-Cerralbo), (Ana Serrano), (Irene Blasco-Machín), (Fadi Hamdan), (Juan Martínez-Oliváan), (Esther Broset), (Pierpaolo Bruscolini).

## Abstract

The design of oligonucleotides with uniform hybridisation temperatures is essential for successful gene synthesis. However, current computational tools for oligonucleotide design face significant limitations, including difficulties in processing long DNA sequences, poor adaptability to specific experimental conditions, limited control over oligonucleotide length, and challenges in minimising spurious dimer formation. To address these issues, we developed CertPrime, an innovative tool designed for scalable and efficient handling of long DNA sequences. CertPrime enables precise customisation of experimental parameters, provides flexibility to limit the maximum oligonucleotide length, and generates designs with reduced deviations in melting temperatures across overlapping regions compared to existing tools. We experimentally compared CertPrime designs with benchmark design methods for a complex DNA sequence and found that CertPrime design led to more efficient gene assembly, significantly reducing the occurrence of non-specific bands. These results make CertPrime a powerful and versatile tool for oligonucleotide design in gene synthesis applications.

**Highlights:**

- CertPrime improves oligonucleotide design by minimising melting-temperature deviations and spurious dimer formation.
- The new tool enhances gene-synthesis efficiency, allowing a precise con- trol over oligonucleotide lengths and experimental parameters.
- Certprime is a scalable tool capable of designing over 99% of the human genome with optimized sequence assembly

**Graphical Abstract:** 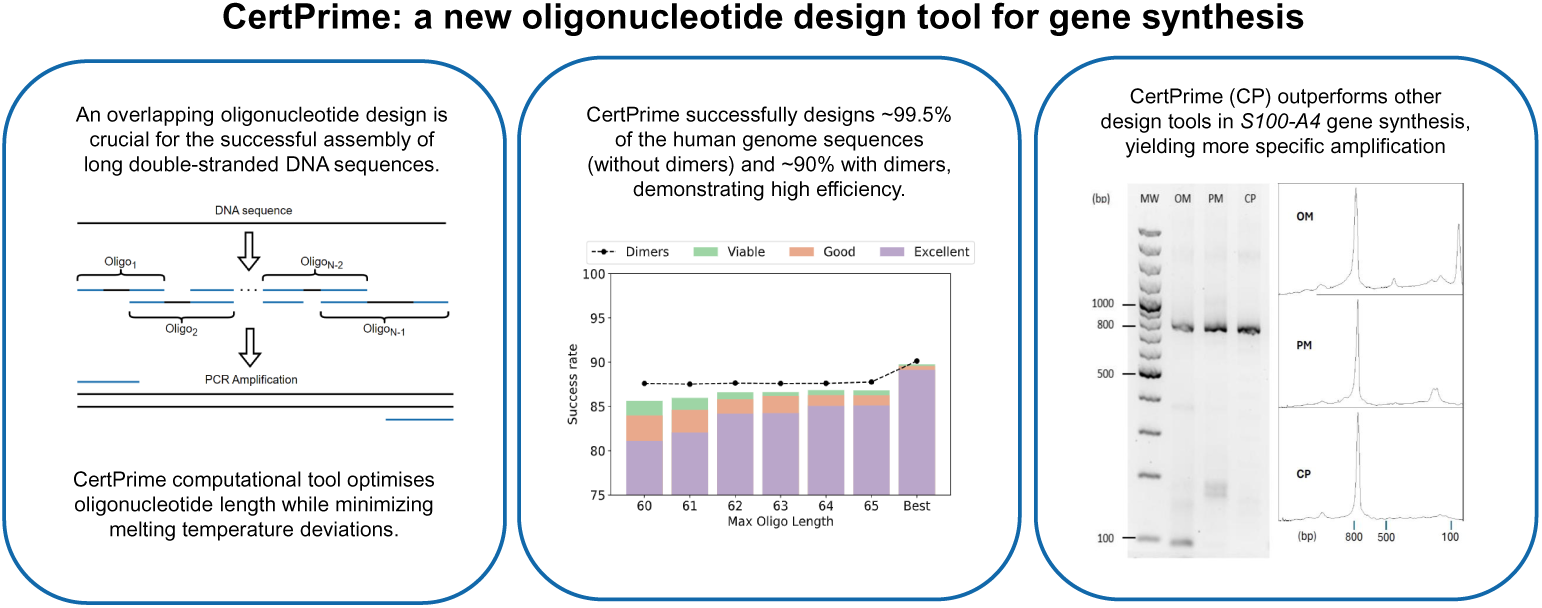

## 1. Introduction

Gene synthesis is a key technique in molecular biology, enabling the generation of long, custom-designed DNA sequences. This capability has driven significant progress in medicine, including the development of gene therapies [1], the production of recombinant proteins for targeted treatments ([2], [3]), or the advancement of mRNA vaccines [4], which have been pivotal in addressing global health challenges such as the COVID-19 pandemic.

The primary method for synthesising genes is based on Polymerase Chain Reaction (PCR) using overlapping oligonucleotides. These short, complementary oligonucleotides are designed with overlapping regions that anneal to adjacent sequences, providing a template for seamless assembly. Through iterative cycles of denaturation, annealing, and extension, DNA polymerase efficiently fills in and joins these overlapping fragments into a continuous DNA sequence. For this process to be effective, it is crucial that the melting temperatures *T_m_* of all the overlapping complementary regions of adjacent oligonucleotides are roughly the same, and align with a target value (*T* ^∗^) pro- vided by the user, which is determined by the optimal operating temperature of the DNA polymerase used in the reaction [5]. This is a necessary (though not sufficient) condition to ensure that all overlapping oligonucleotides de- naturate or anneal at once, yielding a uniform and efficient amplification of all parts of the gene. For this reason, all design tools require an estimation of the melting temperature of the overlapping regions, even if they use this information within different frameworks. The design of overlapping oligonucleotides can be categorised into two primary approaches: gapless PCR and gapped PCR. In the former, the design is such that oligos in the same strand are contiguous, to ensure high-fidelity synthesis. In the latter, neither strand is completely covered by the designed oligonucleotides, that present gaps between them; the complete gene sequence is recovered just because the overlapping oligos, in the opposite strand, span the gaps and provide the missing information. Gapped PCR provides greater flexibility and cost efficiency for specific applications.

Several computational tools and algorithms have been developed, implementing both design strategies. For gapless PCR, tools like TmPrime [6] and DNAWorks [7] are widely used, while Gene2Oligo [8], GeneDesign [9], Assembly PCR OligoMaker [10], Primerize [11], Integrated [12] and DeepFirstSearch [13] are applied for gapped PCR. However, these tools face significant challenges such as difficulties in processing long DNA sequences, poor adaptability to specific experimental conditions, limited control over oligonucleotide length, and challenges in minimising spurious dimer formation.

Regarding difficulties in handling long DNA sequences, tools such as Primerize and Integrated either lack the capability to process extended sequences or experience substantial increases in computational time as sequence length grows, reducing their practicality for large-scale gene synthesis projects. To overcome these difficulties, several experimental methods have further been developed to enable the synthesis of longer DNA sequences through PCR. These include techniques such as thermodynamically balanced inside-out (TBIO) ([14], [15]), dual asymmetrical PCR (DA-PCR) [16], overlap extension PCR (O-PCR) ([17], [18]), PCR-based two-step DNA synthesis (PTDS) ([19], [20]) or PCR-based accurate synthesis (PAS) [21]. While these approaches primarily rely on modifying the PCR experimental process, often by adding additional steps, they do not address the computational challenges associated with oligonucleotide design, leaving a critical gap in scalability and precision for complex DNA assembly projects.

Another critical consideration in oligonucleotide design is the need to constrain the maximum length of oligos. Long oligonucleotides are challenging to synthesise chemically, often resulting in low yields and a higher probability of errors [22]. Tools like Primerize and PCR OligoMaker effectively address this requirement by allowing users to set maximum length constraints. In contrast, algorithms such as DeepFirstSearch and Integrated do not offer this flexibility and often default to designing shorter oligonucleotides, possibly to avoid the challenges associated with longer sequences. However, this bias results in a higher number of oligonucleotides being required to synthesise the same DNA sequence, as compared to tools like Primerize or PCR Oligo- Maker. While this approach does not compromise design quality, handling a larger number of oligonucleotides is less practical in laboratory workflows, increasing complexity and potentially impacting overall efficiency.

Experimental conditions, including DNA concentration, divalent ion levels and the concentration of dNTPs (deoxyribonucleotide triphosphate), can significantly influence oligonucleotide hybridisation temperatures and overall PCR performance [23]. Although PCR OligoMaker and Primerize offer limited options for adjusting DNA and monovalent ion concentrations, and others allow modifications to dNTP and divalent ion levels, these features are often insufficient to comprehensively accommodate the diverse requirements of different experimental setups. This mismatch between design parameters and actual laboratory conditions can lead to discrepancies in melting temperatures and hybridisation efficiency, often resulting in failed PCR assemblies. Consequently, users are frequently left to rely on trial-and-error adjustments, increasing experimental inefficiency and compromising reproducibility.

The formation of spurious dimers is another critical consideration in oligonucleotide design. Homodimers (self-interacting oligonucleotides) and heterodimers formation (unintended interactions between oligonucleotides from different regions of the sequence) often lead to non-specific DNA amplification, reducing the overall yield and accuracy of the synthesis process. While algorithms such as Integrated and DeepFirstSearch incorporate mechanisms to effectively prevent dimer formation, PCR OligoMaker lacks any functionality to address this issue. Similarly, Primerize does not actively prevent dimer formation but offers warnings about their potential presence and identifies the specific oligonucleotides affected, allowing users to manually adjust the design.

To address the limitations of existing design algorithms, we have developed CertPrime, a novel tool designed to overcome these challenges. CertPrime builds upon the principles of the DeepFirstSearch algorithm but re- visits and reformulates each step to enhance performance. CertPrime exhibits excellent computational scalability, enabling it to handle long DNA sequences efficiently. It allows precise customisation of experimental parameters, including the concentrations of divalent and monovalent ions, DNA, and dNTPs, to align closely with laboratory conditions. Additionally, it enables users to set the maximum length for the oligonucleotides, offering flexibility to adapt to the constraints of chemical synthesis.

Notably, CertPrime achieves designs with reduced deviations in the melting temperatures of overlapping regions outperforming PCR OligoMaker and Primerize tools, while maintaining a similar number of oligonucleotides required for synthesis. When tested on the design of a complex DNA sequence with high GC% variations, CertPrime outperformed PCR OligoMaker and Primerize by enabling more efficient experimental gene assembly and significantly reducing the occurrence of non-specific bands. These features make CertPrime a robust and versatile tool for improving the precision, efficiency, and reproducibility of oligonucleotide-based gene synthesis workflows.

## 2. Materials and Methods

### Estimation of the melting temperatures

The melting (or denaturation) temperature *T_m_* of the overlap region of two interacting oligonucleotides, is the fundamental parameter to determine if a design fulfills the desired requirements. *T_m_* is the temperature at which half of the double-stranded structures denature, causing the strands to separate, and it depends non-trivially on the length and on the sequence of the strands, as well as on the characteristics of the solvent. Therefore, accurate estimations of *T_m_* are quite challenging, due to the polymeric nature of the system and to the numerous degrees of freedom involved. The challenge is even harder in the case of non-complementary interacting strands, that one should consider to prevent the formation of spurious dimers with a *T_m_* in the PCR operating range. Finally, the *T_m_* values also depend on the properties of the solution, as the presence of ions or other complex molecules can alter the hybridisation equilibrium and shift the melting temperature. To estimate *T_m_*, most design tools use the nearest-neighbour (NN) thermodynamic model ([24], [25]) that offers a reliable approximation at a reasonable computational cost, when mismatches are absent or isolated along the sequence. This model calculates *T_m_* as the temperature at which the free-energy difference between hybridised and free oligonucleotides becomes zero. The denaturation free energy is derived from the coupling energy of paired nucleotides, the stacking energy of neighbouring pairs and their entropies.

However, when two aligned oligonucleotides present two or more consecutive mismatches, conventional NN models lack the thermodynamic parameters required to accurately predict their effects. Thus, NN models are perfectly fit to calculate the *T_m_* of complementary oligonucleotides, but are unsuitable to calculate the *T_m_* of non-complementary dimers. To overcome this limitation, we took advantage of the temperature-estimation features of the Primer3 software, which implement and enhance the NN model’s capabilities, providing an estimate of the melting temperatures for generic dimers. Primer3 is still based on the NN model, but unlike other tools, it can handle candidate complementary strands of different lengths, since it performs a sequence-alignment, seeking for the best pairing, before estimating the *T_m_*. In case of multiple consecutive mismatches, Primer3 calculates and outputs the *T_m_* of the most stable paired subsequence. Although the predicted *T_m_* may differ significantly from the experimental values, and more sophisticated interpolation schemes could be implemented to improve those predictions, these characteristics of Primer3 prompt for the detection of possible spurious homodimer and heterodimer interactions, since, in practice, the size of the overlap regions limits the magnitude of the estimation error.

Additionally, Primer3 allows users to specify the values of experimental parameters, such as the concentrations of monovalent and divalent ions, the concentration of dNTPs, as well as the DNA concentration. This flexibility enables a more accurate reproduction of experimental conditions, thereby improving the reliability of oligonucleotide design and minimising the risk of unintended interactions during synthesis.

### Computational protocol for oligonucleotide design

Our method builds upon and enhances the Integrated strategy [12] that consist of four steps. In the first one, the DNA is divided into fragments with similar melting temperatures (of the dsDNA), all close to the target temperature *T* ^∗^ specified by the user. Notice that it is sufficient to consider a single strand of DNA, since the complementary sequence is completely determined. The above partition is done using a greedy algorithm that selects the (locally) best solution, starting from one end of the DNA and proceeding towards the other. The fragments thereby obtained represent the candidate regions from which the overlap regions will be extracted.

The second step involves an iterative process, analogous to a zero-temperature Monte Carlo algorithm. During this phase, the boundaries between segments are adjusted by shifting them by 0 to 4 nucleotides. These adjustments aim to reduce fluctuations in the *T_m_* values of the segments, because the greedy algorithm does not guarantee a globally optimal outcome. Subsequently, between 0 and 5 nucleotides are removed from both the 5’ and 3’ ends of each segment, resulting in 36 candidate overlap regions for each segment. Finally, a dynamic programming algorithm is employed to select the optimal overlap region candidate for each segment. This approach systematically explores all possible combinations of the trimmed ends, identifying the overlap regions that minimise the melting-temperature deviations and enhance the overall efficiency of the assembly process.

This process is efficient but has several drawbacks. Specifically:

- The greedy algorithm employed in the first step yields a locally optimal outcome, but not a globally optimal one. When approaching the end of the strand, there is little control over the *T_m_* of the last segment, which is often significantly lower than the others, potentially causing difficulties in the assembly process.
- The subsequent iterative algorithm, that redistributes the positions of the ends of the fragments, only accepts changes that improve the overall homogeneity of the *T_m_*. However, all these temperatures, except for that of the last fragment, have already been adjusted to a local optimum in the previous stage. As a result, the initial iterations only shift the boundary between the last two segments; then, the other boundaries are moved to try to redistribute the residual mismatch between the actual and the target temperatures. This results in the melting temperature of the fragments being overall slightly lower than the target temperature specified by the user.
- The dynamic algorithm imposes no constraints on oligonucleotides length, prioritising segments with *T_m_* closest to the target value. However, since the iterative algorithm produces slightly lower *T_m_* than the desired value and base removal at the third step of the algorithm further decreases the *T_m_*’s, the base removal is actually discouraged, which results in almost gapless designs.

To cope with these issues, we began by modifying the initial greedy algorithm, to yield fragments with higher melting temperatures, by setting the initial target temperature to a *T* ^′^ *> T* ^∗^, depending on the fraction of GC nucleotides in the strand. In the new procedure we also check, and enforce, that the segments’ lengths fall in the range [*l^ov^_min_* and *l^ov^_max_*] between the minimum and maximum overlap lengths, specified by the user. Since the *T_m_* of a pair of hybridised strands is monotonically increasing with their length, the temperature *T_m_* is always set higher than the target *T* ^∗^ provided by the user. This ensures that shortening the regions *S_i_* will bring their melting temperature *T_m,i_* closer to the desired target (Figure 1).

**Figure 1:**
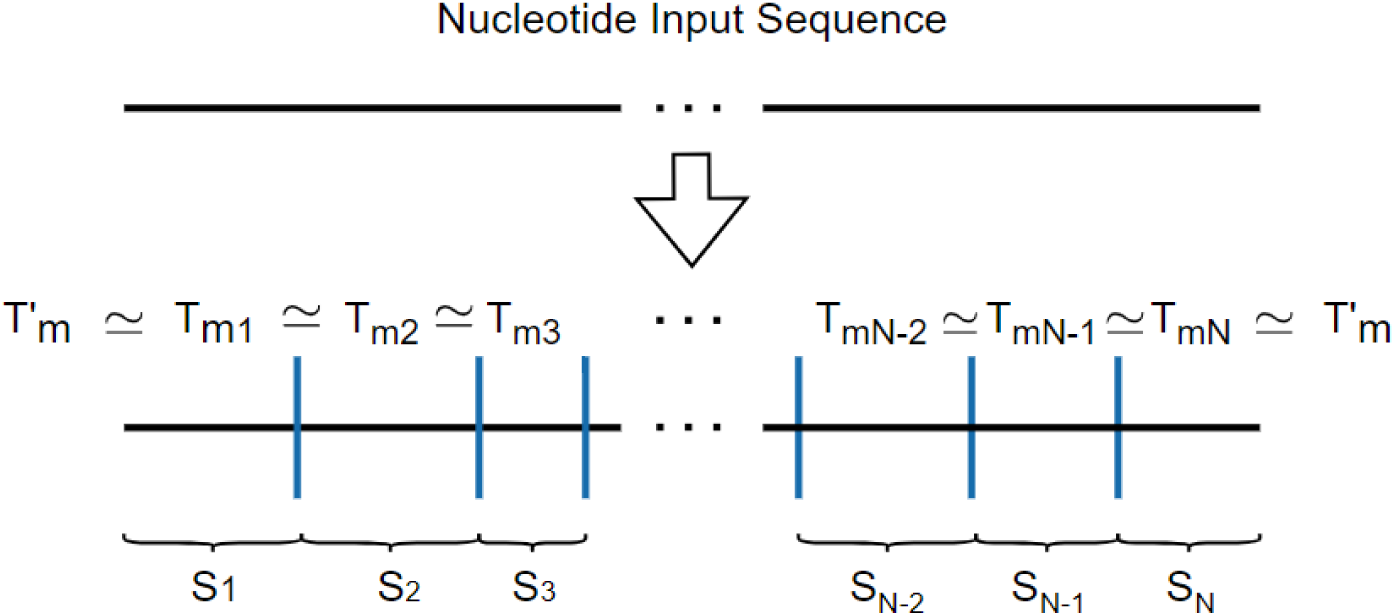
Scheme of the first step in the design of CertPrime oligos. In this process, we start with an input sequence and obtain N segments, each with similar *T_m_*, close to an increased target temperature *T* ^′^ . The number of segments *N* is forced to be odd, so that the final number of oligonucleotides, *N −* 1, is even.

Furthermore, we modified the greedy algorithm to prevent edge effects in the final fragments. Specifically, when we reach a position located between three and four *l^ov^* from the end of the sequence, we determine the last three or four segments altogether. The exact number of segments depends on the fragments already set, and is determined by the need to have an odd total number of fragments (and hence, of overlap regions), ensuring that the final number of oligonucleotides is even. This constraint is related to technical reasons, to cope with the experimental setup and reconstruct the dsDNA from the oligo-design in a single PCR run, using an excess concentration of the terminal oligonucleotides. An even number of oligonucleotides in the design ensures that the first and last ones are located on opposite strands. This arrangement allows the entire sequence to be duplicated effectively.

In the second step of the Integrated approach, the iterative process is replaced by a Simulated Annealing (SA) method to displace the fragment ends, and obtain a more uniform distribution of melting temperatures around the target value *T* ^′^_m_ . Additionally, throughout the SA process, we ensure that the lengths of the segments remain within the specified bounds, *l^ov^_min_* and *l^ov^_max_*. At this point, the outcome is a gapless design.

In the following step, gaps are introduced to improve the overall quality of the design. As in the Integrated approach, we remove nucleotides from both the 5’ and 3’ ends of each segment, but without restricting the removal to just 5 nucleotides. Instead, we generate all possible fragments that satisfy the constraints on the overlap length (Figure 2).

**Figure 2:**
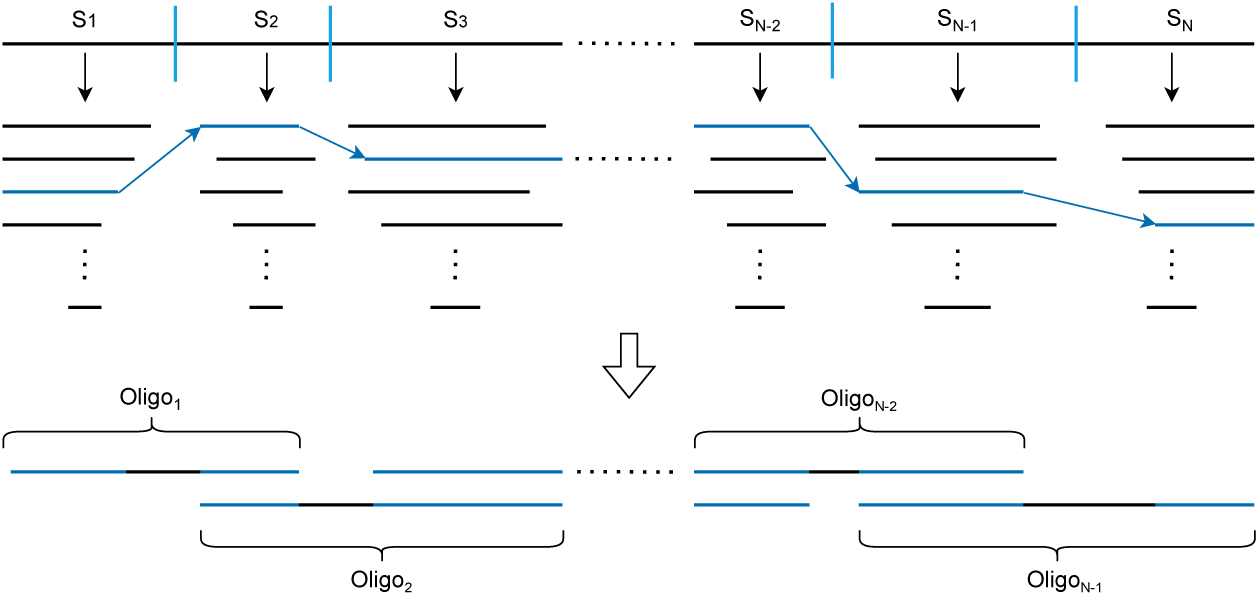
Schematic representation of the final step performed by CertPrime. The process starts with the N fragments obtained with the gap-free partition described in Figure 1. Candidate overlap regions (black segments) are obtained upon removing *i* nucleotides from the ends of the *S_j_* fragments. The choice of these segments (blue lines) is performed to minimise Equation 2 while keeping all the final oligonucleotide lengths below *M.O.L*.

Finally, a selection of the candidate overlap regions among the latter fragments is made, which produces the final oligonucleotide design, according to the following criteria:

- The length of the selected fragments fall inside the user-specified range

for overlap regions [*l^ov^_min,_ l^ov^_max_*], and the oligonucleotides spanned by any pair of neighbouring fragments are shorter than the maximal oligonucleotide length *M.O.L*, specified by the user.

- The overall deviation of their melting temperatures *T ^i^* from *T* ^∗^ is minimised.

The length criterion above is a hard constraint, stronger than the requirement on the melting temperatures, ensuring that the resulting lengths of the oligonucleotides and overlap regions always fall within the specified ranges, irrespective of temperature deviations. Based on these criteria, we apply a dynamic programming algorithm that finds the best design from the set of fragments generated in the previous step. This approach minimises the deviations in *T_m_* while assigning an infinite deviation to solutions that do not meet the required oligo-length criterion. Once the design is complete, a few steps are still necessary to check the results and translate them into a proper output for the user:

- The strand of the even-numbered oligonucleotides is changed, returning their reverse complementary sequence.
- The presence of homodimers and heterodimers is checked, by calculating the melting temperatures for each possible interaction between pairs of oligonucleotides in the design.
- The overlap regions are explicitly extracted, and their melting temperature is determined.
- The %GC content for both the overlap regions and the oligos is calculated.

The input parameters that must be specified by the user, along with the gene sequence, are reported in Table 1, together with the values used in all the computational tests in this article.

**Table 1:** Input parameters. List of user-specified input parameters for CertPrime. The reported values are those used in the computational tests. *T* ^∗^ is the target melting temperature for the overlap regions, M.O.L. is the maximum length for any oligonucleotide in the design, [*l^ov^_min,_ l^ov^_max_*] specifes the range of possible lengths for the overlap region. S.O. indicates short oligos, and m.O.L. (S.O.) is their minimum length. Brackets indicate concentrations; dNTPs: deoxyribonucleotide triphosphates.

|  |  |  |  |  |  |
| --- | --- | --- | --- | --- | --- |
| [Na<sup>+</sup>] | 50 mM | [dNTPs] | 0.6 mM | $(l_{min}^{ov}, l_{max}^{ov})$ | (16-30) bp |
| [Mg<sup>+2</sup>] | 1.5 mM | $T_m^*$ | 60 °C | $T_m^*$ (S.O.) | 60 °C |
| [DNA] | 300 nM | M.O.L. | 60-65 bp | m.O.L. (S.O.) | 17 bp |

### 2.1. Short-primers design

The PCR experimental protocol relies on two distinct primers (the sense and antisense one) that define the region to be amplified by annealing to the 3’ ends of the template and complementary strands. Thus, in order to amplify the whole gene and prevent partial or incomplete amplification, it is important that the primers anneal to the first and last oligonucleotides of the design with the highest probability. Instead of simply using an excess concentration of the terminal oligonucleotides, an alternative effective strategy involves designing ’short’ oligo primers, with a melting temperatures of the oligo as a whole (*T_m_*(S.O.)) close to a target value (*T* ^∗^ (S.O.)), and a given minimum length (m.O.L.(S.O.)). These short primers are complementary to part of the first and last oligonucleotides in the design, thereby enhancing both efficiency and specificity.

CertPrime allows to design such short oligos, using the following protocol: it takes the first or last *k* bases, with *k* equal to the minimum oligo length for short oligonucleotides (m.O.L.(S.O.)), from either the initial or final oligo, depending on which S.O. is being designed. It then continues adding bases until the distance between the melting temperature of the short oligo (*T_m_*(S.O.)) and the target value is minimised.

### 2.2. Method-assessment protocol

#### 2.2.1. Human genome database preparation

To perform an *in-silico* evaluation of the performance of the method, the ‘Homo Sapiens’ database was downloaded from Ensemblr [26] on 10 April 2024. This database contains 122,676 sequences, with lengths ranging from 10 to 10,976 nucleotides (Figure S.I. 1). Two subsets were extracted from the database: one for the synthesis-quality test (DS1) and another for the computational-scaling test (DS2). For the former, sequences were filtered to encompass lengths ranging from 150 to 1350 nts, resulting in a dataset of 73.424 sequences, to be used to assess how many human genes can be reliably designed by CertPrime, For the latter, a dataset of 960 sequences was built, sampling 10 sequences from the human database for each length, in multiples of 50, ranging from length *L* = 250 to *L* = 5000 nucleotides. This dataset was used to test the dependence of the computational cost on sequence length.

#### 2.2.2. Indicators of design quality

For each design, we calculate the following quality indicators:

- The temperature range:

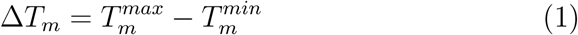

- The distance from target:

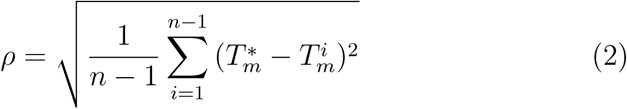

- The temperature spread:

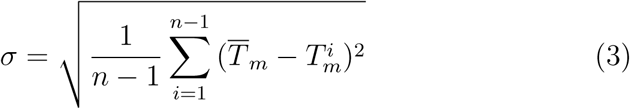

Here, *n* represents the number of oligonucleotides in the design; *T ^i^* denotes the melting temperature of the overlap (hybridisation) region between oligonucleotides *i* and *i* + 1; 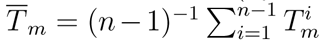 is their average melting temperature, while *T* ^min^, *T* ^max^ correspond to the minimum and maximum melting temperatures, respectively, appearing in the oligo design. Additionally, *T* ^∗^ represents the desired (user-specified) melting temperature for the hybridisation regions.

The first indicator, Eq. 1 highlights designs presenting hybridisations with very different melting temperatures, that could affect the assembly process. Equation 2 reports on the overall distance of the *T_m_*values from the tar- get melting temperature *T* ^∗^, ensuring consistency of the design with the desired experimental conditions. Finally, the standard deviation Eq. 3 measures how closely the different *T_m_* values are clustered together, providing an assessment, complementary to Eq. 1, of uniformity across all overlaps.

The combined use of multiple indicators, rather than relying solely on the standard deviation Eq. 3, as done in other studies (e.g. Integrated, DeepFirstSearch or TmPrime), allows us to identify and mitigate potential issues related to melting temperature variability, thereby enhancing the efficiency and reliability of the gene synthesis process.

Based on the values of these three indicators, we proposed a classification of the designs as:

- “Excellent” (*ρ <* 1.5 and *σ <* 1 and Δ*T_m_ <* 2)
- “Good” (*ρ <* 2.5 and *σ <* 2 and Δ*T_m_ <* 3)
- “Viable” (*ρ <* 3.5 and *σ <* 3 and Δ*T_m_ <* 4)

In this way, we can effectively evaluate and select oligonucleotide configurations that are more suitable for successful gene synthesis.

In addition to these three indicators, we also check the potential formation of dimers, to avoid spurious interactions in the design. This will serve as another criterion for evaluating oligonucleotide designs, ensuring that unintended homodimer and heterodimer formations are minimised, to enhance the efficiency and specificity of the gene synthesis process.

#### 2.2.3. Synthesis of DNA sequences

A mix of desalted oligodeoxynucleotides (Integrated DNA Technologies) was prepared in TE buffer 1X, with a ratio outer/inner oligos of 40. The first PCR reaction (gene assembly) was performed in a final volume of 25 *µ*L, with 40 pmol of each outer oligo and 1 pmol of each inner oligo, 100 mM KCl, 3% DMSOI and 1X PrimeSTAR Max Premix (Takara). The reaction mixture was subjected to PCR amplification under the following conditions: initial denaturation at 98°C for 30 s, followed by 20 cycles of denaturation at 98°C for 5 s, annealing at 60°C for 10 s, and extension at 72°C for 30 s. A final extension step at 72°C for 5 minutes was performed after the last cycle. For the second PCR reaction (gene amplification), 2 *µ*L of the crude product from the first PCR was mixed with 40 pmol of each outer oligo, 100 mM KCl, 3% DMSO and 1X PrimeSTAR Max Premix (Takara), and double-distilled water to a final volume of 25*µ*L. The thermal cycling conditions were: initial denaturation at 98°C for 30 s; 30 cycles of denaturation at 98°C for 5 s, annealing at 60°C for 10 s, and extension at 72°C for 30 s; followed by a final extension at 72°C for 5 minutes. PCR products were purified using QIAquick PCR Purification Kit and then analysed using 1% agarose gel electrophoresis.

## 3. Results

### 3.1. Performance of CertPrime in designing human genes

We generated oligonucleotide designs for all the sequences in the data set DS1, using the parameters reported in Table 1, with a target melting temperature *T* ^∗^ = 60 ^◦^C. To explore the effect of oligonucleotide length on design efficiency and performance, we considered constraints on the maximal oligonucleotide length in the design ranging from 60 to 65 nucleotides. This approach resulted in six different designs for each sequence, allowing us to assess the versatility and robustness of CertPrime across various length constraints. Then, the designs were classified as “Excellent”, “Good” or “Viable” as specified in the Methods section. The results, reported in Figure 3, suggest that increasing the maximum length of the oligonucleotides enables us to design a higher percentage of the genome satisfactorily, as CertPrime gains more flexibility in the design process. Specifically, satisfactory designs were achieved for more than 97% of the genome across all maximum lengths tested, this percentage exceeding 99% when the maximum length was set to 65 base pairs. Lastly, when choosing the best design for each sequence irrespectively of the maximal length, we were able to design 99.6% of the genome at “Excellent” level. Notice that the best design for a certain maximal oligo length is not necessarily found when designing at bigger lengths, due to the stochastic nature intrinsic to the design process. This highlights CertPrime’s exceptional adaptability to the diversity of sequences found across the entire human genome.

**Figure 3:**
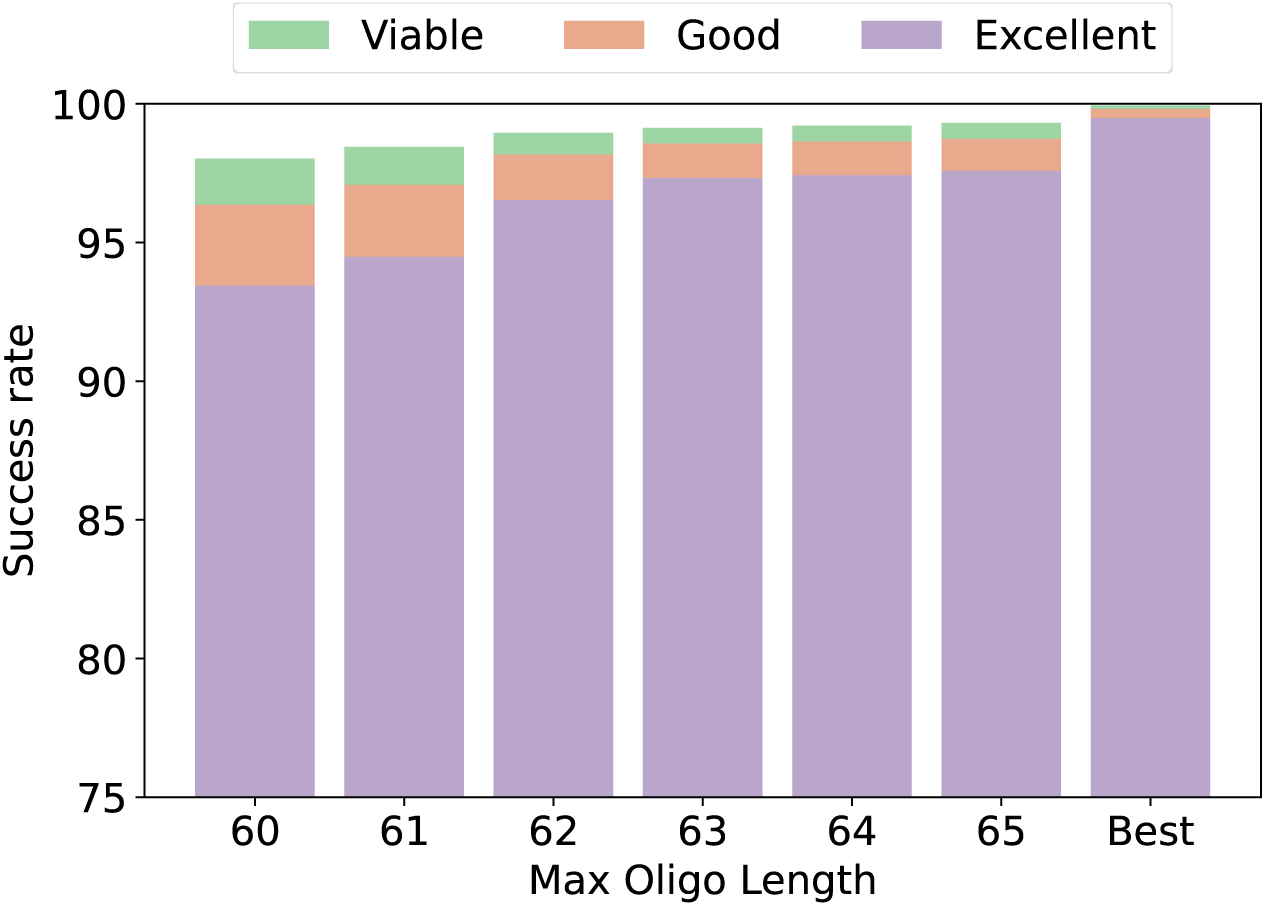
T**h**e **success rate and classification quality for designs with different constraints on the maximum oligo length**. The choice of the lengths is related to the typical values managed in the experimental setups. The column “best” reports the best design for each sequence, irrespective of the maximum oligo length for which it was obtained.

To further evaluate CertPrime, we applied a “post-filtering” step to refine the designs by accounting for dimer formation imposing that all designs that are affected by homodimer or heterodimer pairings with a *T_m_* greater than (*T* ^∗^ *−* 5)^◦^*C*, are rejected, independently from how well they performed in terms of *σ* or Δ*T_m_*.

Using this approach, we obtained the black dotted line in Figure 4. The results indicate that the dimers rendered a fraction between 12% and 14% of the designs invalid, depending on the maximum oligo length. Furthermore, as shown in the column “Best”, we observed that in 9% of cases, no valid design was identified. This suggests the existence of “dimer-prone” sequences, for which it is impossible to achieve dimer-free design even when allowing some flexibility on the oligonucleotide lengths.

**Figure 4:**
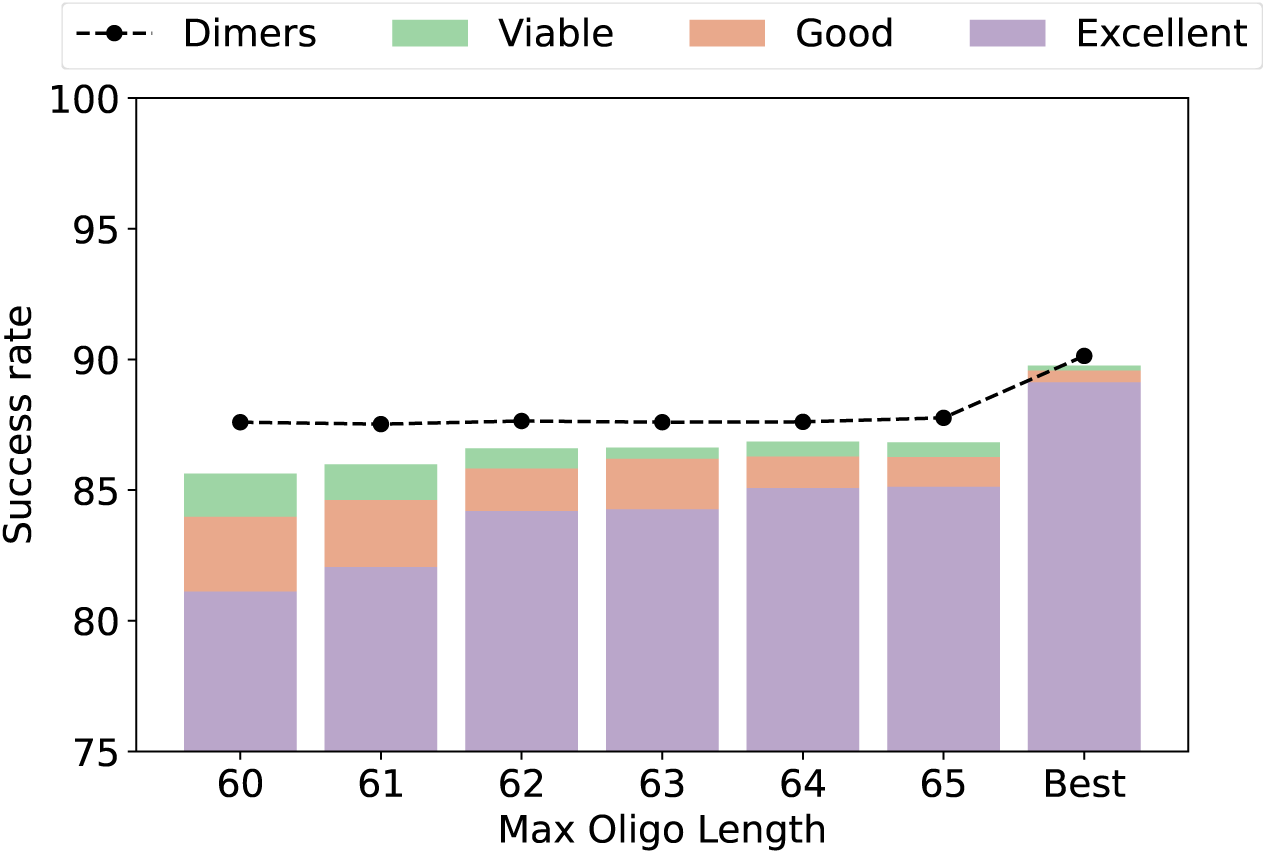
C**l**assification **of the same designs as in Figure 3 but discarding ho- modimers and heterodimers.** The black dotted line “Dimers” marks the maximum possible success rate after discarding designs with dimers with a *T_m_* of more than 55^◦^*C*. In the “Best” column, the best design has been selected for each sequence.The gap be- tween the colored bars and the corresponding black dot represents sequences that are not dimer-prone, but for which no satisfactory design could be found.

Results also suggest that, especially for stronger limits on the maximum oligo-length, there were sequences that in principle were not affected by dimers, but for which CertPrime could not find any good design, according to the *σ* or Δ*T_m_* criteria.

Next we determined whether the percentage of invalid designs, caused by either dimer formation or CertPrime constraints, varies with sequence length, with the aim to identify if there is an optimal sequence length that maximizes the probability of a successful design.

For statistically reliable results, we considered the same database DS1 as before, and grouped the sequences, according to their lengths, in 21 blocks, spanning 50 lengths each (i.e. the first block contained sequences of 150-199 nts, and so on until the last block, with 1250-1299 nts).

Results showed that the probability of success (with at least one valid design) starts at values above 92% and it reaches 95% (Figure 5, “Best”). Additionally, as sequence length increases, the probability of success steadily declines. This trend is primarily due to the increase in the number of oligonucleotides required for synthesising longer sequences, which raises the probability of unwanted dimers formation. This suggests that dividing a long sequence into shorter segments and conducting two separate syntheses could be an effective strategy to enhance the quality of the design by minimising dimer formation. This approach can potentially improve experimental yield and overall efficiency.

**Figure 5:**
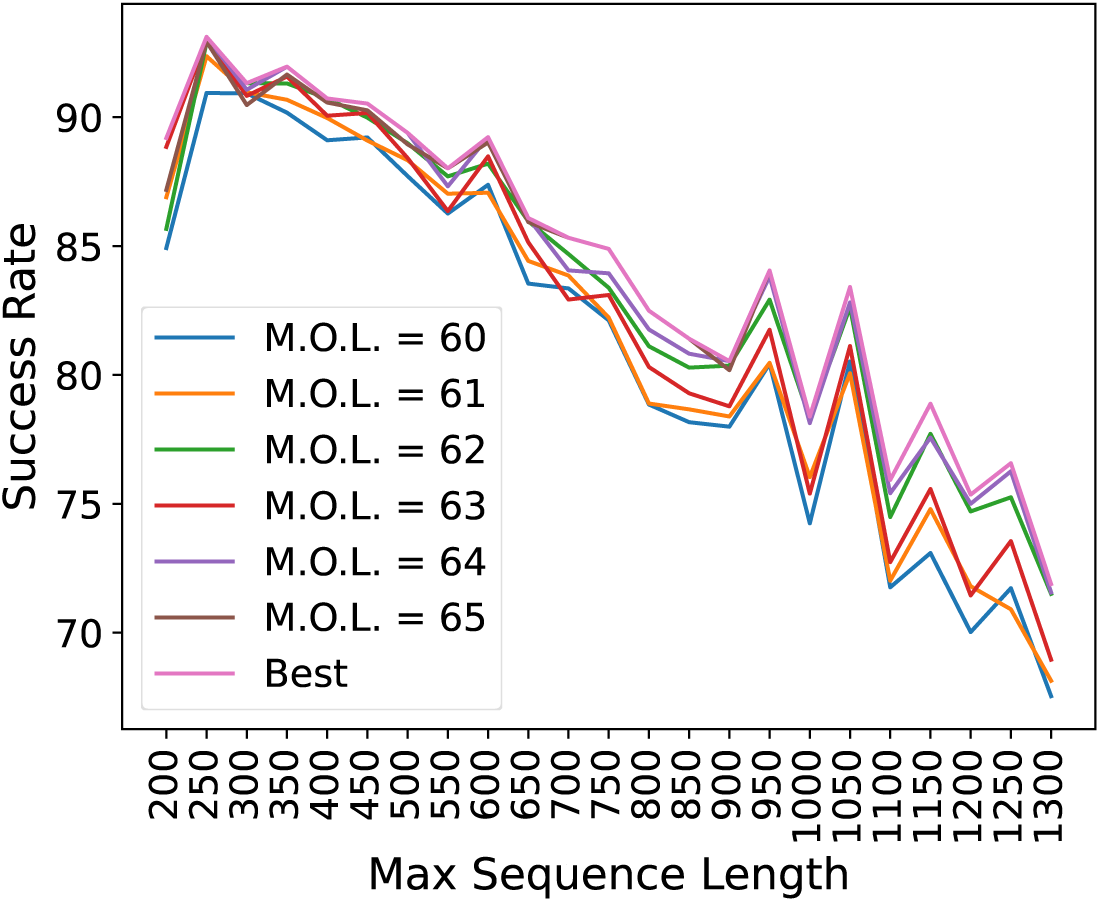
C**e**rtPrime **success rate according to sequence length.** The X axis labelled as “Max Sequence Length” refers to the maximum sequence length in the block under consideration. The “Best” line considers the best design selected for each sequence, regardless of the maximum oligo length (M.O.L.) between 60 and 65 nts.

### 3.2. Computational cost dependence on DNA length

To study the length-dependence of the computational cost of CertPrime, we measured the time required to design each sequence contained in the human genome DS2 (Figure 6). The results suggest a roughly quadratic relationship between the sequence length and the computational time, both when considering and measuring the time required for the process.

**Figure 6:**
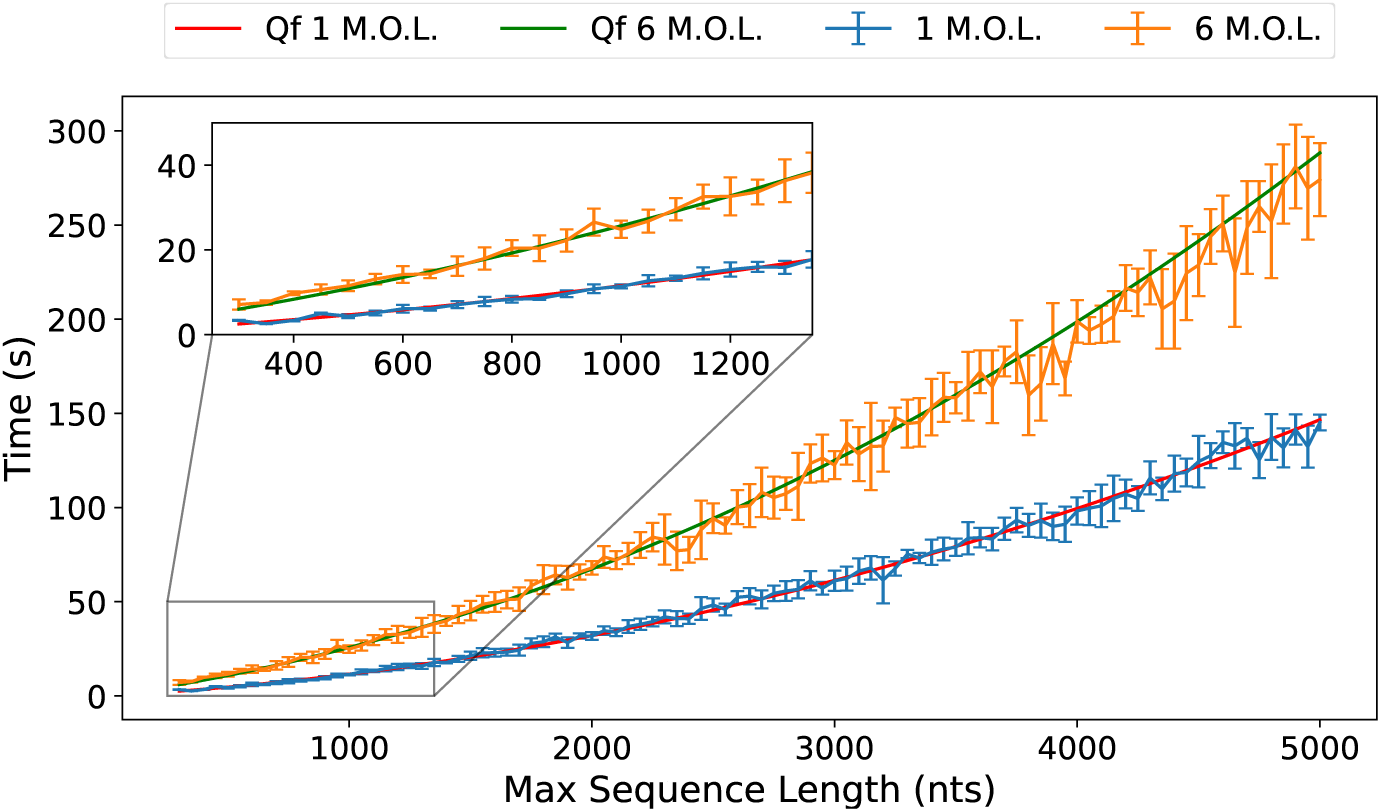
D**e**pendence **of computation time on DNA sequence length.** In the case of the “1 M.O.L.” line (in blue) only one design *per* sequence was generated, with a maximum oligo length (M.O.L.) of 65 nts, while for the “6 M.O.L.” line (in orange) six designs were made with maximum oligo lengths from 60 to 65 nts for each of the sequences. In addition, a quadratic fit (*Q_f_*) was performed for each of the lines, yielding the red and green curves. The tests were performed on a computer with a 24-core Intel i9-13900K processor and 64 Gb RAM.

The results, reported in Figure 6, suggest a roughly quadratic relationship between the sequence length and the computational time, both when consid- ering a single design *per* sequence, with a specific maximum oligo length, or designs across all 6 lengths. For the single length case, we get:

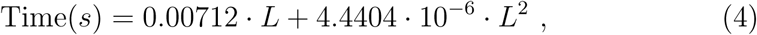

while for the case of the 6 different maximum oligonucleotide lengths:

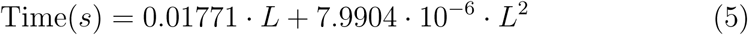

where *L* is the length of the sequence.

From these equations, it can be seen that the computational cost associated with considering several different cases for the maximal oligo length does not scale linearly with the number of cases. This non-linear behaviour arises because CertPrime is parallelised, to deal with each case independently. Consequently, all designs are performed simultaneously, with a significant reduction of the computational cost of the method. This parallelisation strategy enables CertPrime to handle long sequences in a reasonable time, approximately 140 seconds for lengths of 5000 nts.

### 3.3. Assembly efficiency

To assess CertPrime’s ability to generate experimentally viable designs, we used it to create optimal oligonucleotide sets for eight different DNA sequences and evaluated the quality of the designs using the indicators introduced in Eqs. 1, 2, and 3. Table 2 shows that the Δ*T_m_*, *ρ*, and *σ* values are all close to 0, which indicates not only that all temperatures are clustered, but also that they are clustered around the target value. The only sequence that departs from this trend is Seq 1, which gives higher values for The designed oligos were then synthesised and subsequently evaluated in PCR amplification experiments. The quality of the resulting assemblies was then assessed through agarose gel electrophoresis, providing a visual representation of the efficiency and accuracy of the assembly process (Figure 7). We observed that for all sequences, a single intense band corresponding to the expected size of the synthesised fragments was obtained. This indicates that the synthesis was successful and that no dimers or other undesired by- products were formed, including in the case of sequence Seq 1.

**Table 2:**
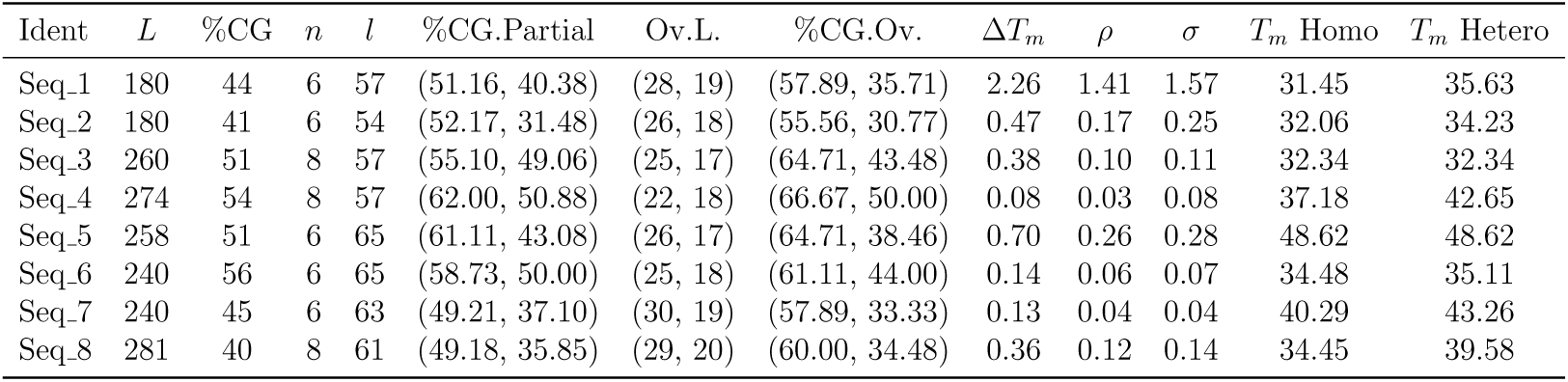
Characteristics of the sequences used for computational and exper- imental tests, and parameters of the corresponding oligonucleotide designs produced by CertPrime. *L* is the sequence length, %GC is the GC content, *n* is the number of oligonucleotides, *l* is the length of the longest oligonucleotide. %CG.Partial represents the GC content of the 50-nucleotide region in the sequence, with the highest and lowest GC content. The column Ov.L. reports the maximum and minimum lengths of the overlap regions, while %GC.Ov reports the maximum and minimum GC percent- ages of the overlap regions. The indicators Δ*T_m_*, *ρ*, and *σ* were introduced in Eq. 1, 2, and 3, respectively. *T_m_* Homo is the melting temperature of the oligonucleotide with the strongest self-interaction (thus forming the most stable homodimer), and *T_m_*Hetero denotes the melting temperature of the pair of oligonucleotides that interact most strongly with each other (forming the most stable heterodimer).

**Figure 7:**
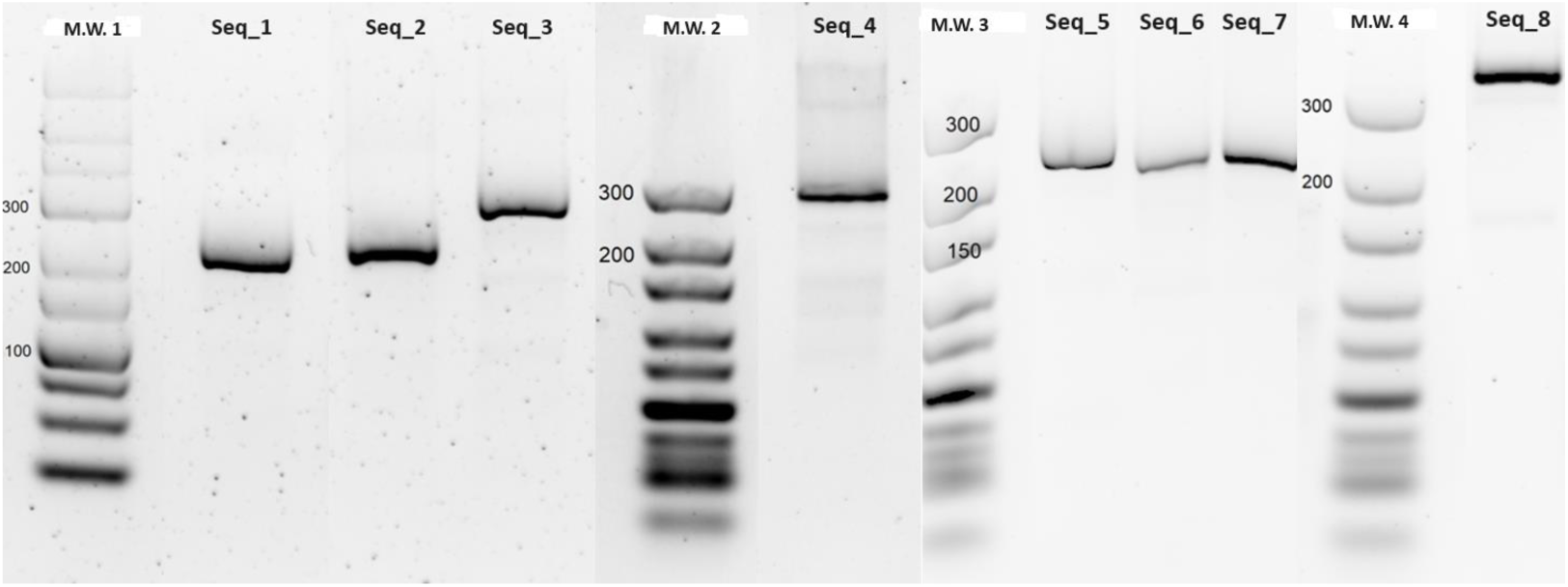
Agarose gel electrophoresis results of the oligonucleotide designs presented in Table 2. The molecular weight (M.W.) lanes correspond to all the sequences to their right until the next M.W. lane.

### 3.4. Comparison of CertPrime with other software tools

The performance of CertPrime was compared to that of other available design programs to evaluate its potential experimental advantages. Specifically, we designed oligonucleotides for the S100A4, PKB2, and GFPuv genes, which are frequently referenced in the literature as benchmarks for evaluating oligonucleotide design tools in gene synthesis studies and which designs are available ([6], [7], [8]). Also, their nucleotide sequences were obtained from [6] and can be found in the Supplementary Information. Thus, we could compare CertPrime’s designs with those of Primerize, PCR Oligo- Maker, TmPrime, DeepFirstSearch, Integrated DNAWorks, and Gene2Oligo (Table 3).

**Table 3:**
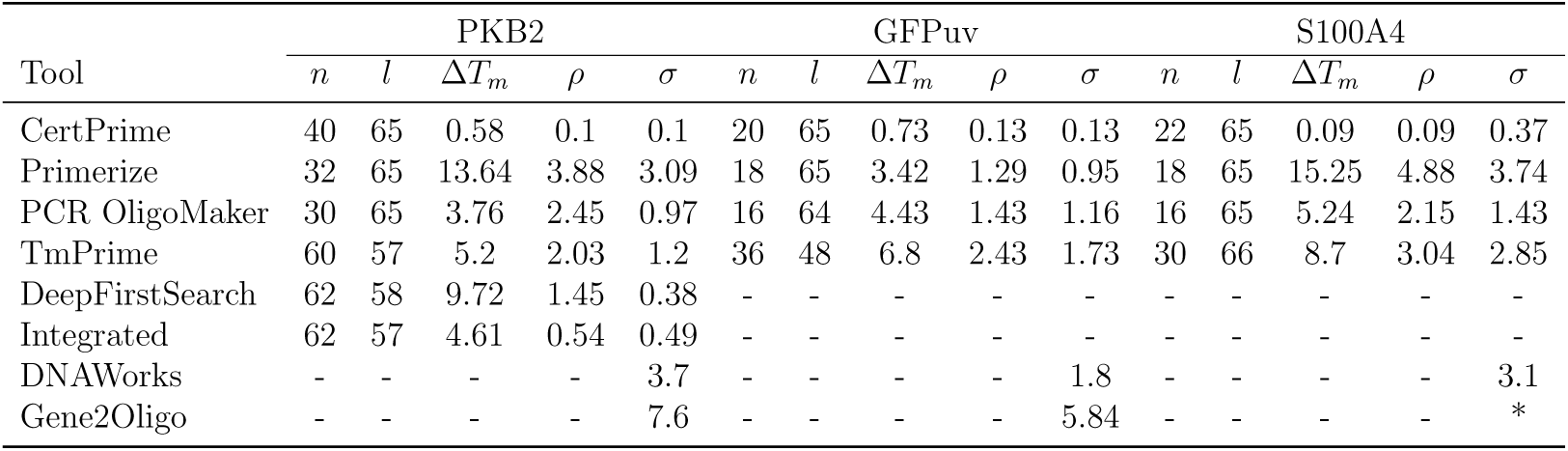
Comparison between designs by different methods for PKB2, GFPuv and S100A4 sequences. Here *n* is the number of oligos in the design, *l* is the length of the longest oligo, and Δ*T_m_*, *ρ* and *σ* are the indicators introduced in Eq. 1, 2 and 3, respectively. For all designs performed with CertPrime, Primerize and PCROligoMaker, the maximum allowed oligo length was set to 65 nts and *T* ^∗^ = 60^◦^*C*. The TmPrime, DNAWorks and Gene2Oligo designs were taken from the TMPrime article [6] where they were found to have *T* ^∗^ = 65^◦^*C* and DeepFirstSearch and Integrated were taken from [13] where the PKB2 and S100A4 *T* ^∗^ = 65^◦^*C* while the GFPuv design had *T* ^∗^ = 55^◦^*C*.

Gene2Oligo and DNAWorks were only partially assessed as their web servers appear as unavailable, and their source code is not publicly accessible. As a result, we report only the *σ* values extracted from their respective publications ([7], [8]). Similarly, DeepFirstSearch and Integrated lack public code or web servers, so parameters could only be retrieved for PKB2 from oligo designs provided in their related publications ([12], [13]). Primerize and PCR OligoMaker were accessible but presented notable limitations. Primerize’s computational scalability restricts input sequences longer than 1000 bp on its web server, although users can bypass this restriction by locally installing the software. The web server of PCR OligoMaker, on the other hand, is not actively mantained, and it is only compatible with outdated browser versions; also, its source code is not publicly available. Despite these constraints, we successfully installed Primerize and accessed the PCR Oligo- Maker web server, and we could produce designs for the S100A4, PKB2, and GFPuv genes with a maximum oligo length of 65 nucleotides and a *T_m_* set to 60°C.

In terms of computational design quality, CertPrime consistently exhibited the lowest values for Δ*T_m_*, *ρ* and *σ* across all tested genes (Table 3), serving as a robust indicator of the method’s accuracy and reliability. Regarding the number of oligonucleotides required *per* design, Integrated, DeepFirstSearch, and TmPrime require more oligos compared to the other methods, whereas PCR OligoMaker and Primerize demonstrate a more efficient use of oligos.

Notably, Primerize, DeepFirstSearch, and Integrated showed significantly higher Δ*T_m_* values compared to their respective *σ* values. This suggests a scenario where all melting temperatures are well-clustered, with the exception of a few outliers that distort the overall design quality. This discrepancy poses a practical challenge in synthesis, as such deviations often lead to experimental failures. These issues arise because these methods focus solely on minimising *σ* without accounting for the behavior of Δ*T_m_*, allowing such errors to remain undetected.

The S100A4 gene presents a region with a very low GC% content, making its design more challenging than that of the PKB2 and GFPuv genes (Figure 2 in S.I.). Therefore, we performed experimental tests on S100A4 gene designs generated by PCR OligoMaker, Primerize, and CertPrime. The designs produced by PCR OligoMaker and Primerize exhibited dimer formation, as evidenced by bands around the 100 base pairs region on the agarose gel electrophoresis (Figure 8, right panel). These dimers were further evidenced through chromatograms corresponding to the gel image analysis (Figure 8, left panel). In contrast, the design produced by CertPrime resulted in a single, more intense band with no detectable dimers at lower mass. This outcome indicates a successful synthesis and assembly process, underscoring CertPrime’s effectiveness in preventing spurious interactions and producing high-quality oligonucleotide assemblies.

**Figure 8:**
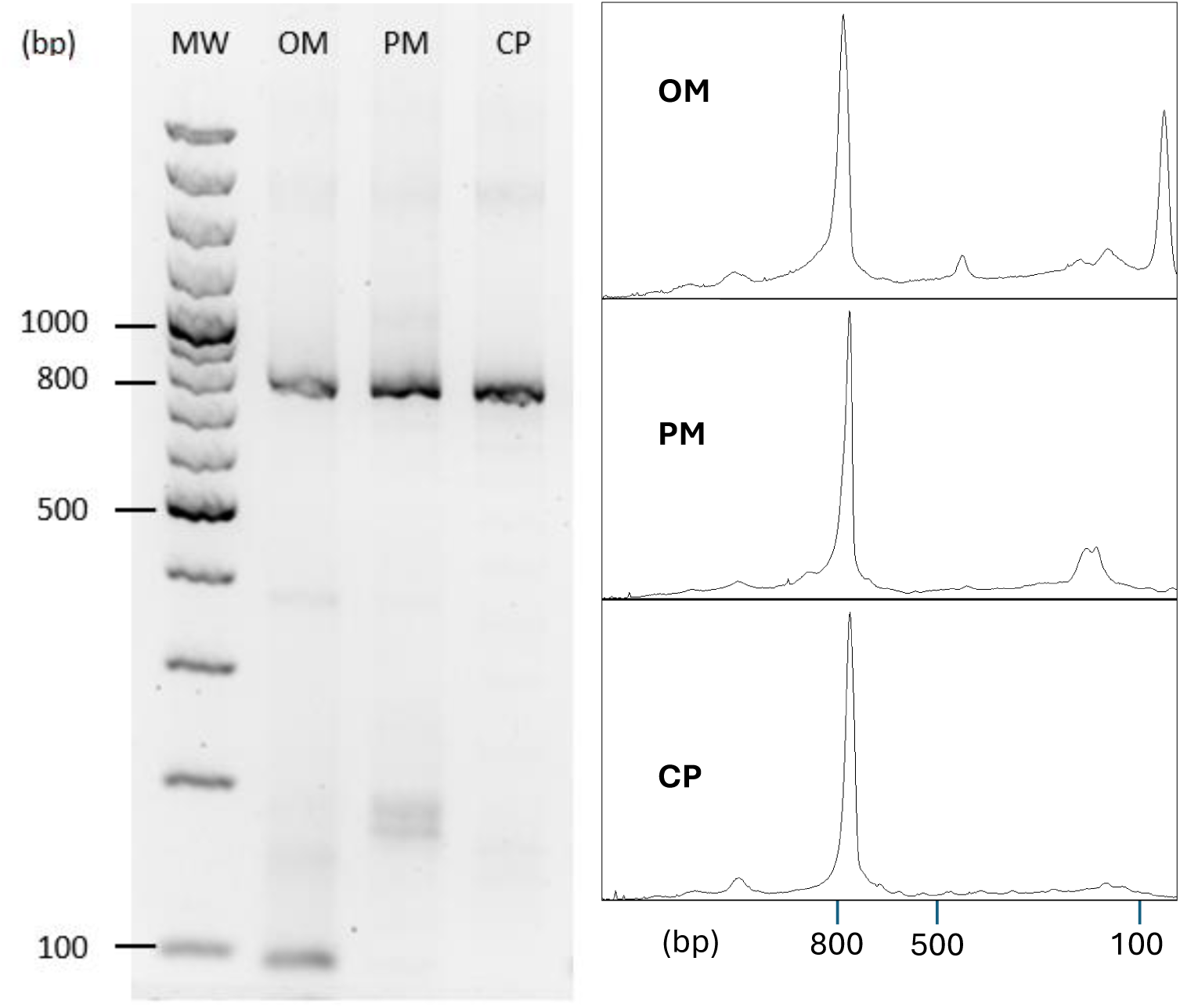
Analysis of PCR amplification of the S100A4 gene using three different primer designs: PCR OligoMaker (OM) Primarize (PM) and CertPrime (CP). The left panel displays agarose gel electrophoresis results for each PCR experiment. The right panel shows signal intensity profiles derived from the corresponding gels, where the intensity of each band correlates with the height of the peak at the respective molecular weight. An optimal amplification is represented by a sharp, narrow peak aligned with the expected molecular weight of the S100A4 gene sequence.

## 4. Conclusions

We have developed CertPrime as a fast, flexible, and highly accurate tool for oligonucleotide design in gene synthesis, with a particular strength in minimising *T_m_* dispersion. By integrating key innovations at every step of the oligo design process, CertPrime achieves substantial improvements in computational efficiency while significantly increasing experimental success rates. One of CertPrime’s prominent strengths lies in its flexibility, enabling precise specification of experimental conditions, that surpasses the capabilities of other available tools. It offers a range of customizable parameters to guide the design process, such as the maximum oligo length and overlap length. Additionally, CertPrime includes an advanced feature to minimise spurious oligonucleotide interactions, further improving the reliability of its designs.

The tool’s performance is underscored by its ability to synthesise over 99.5% of the human genome when dimer formation is not considered, and approximately 90% when dimers are taken into account. Computationally, CertPrime demonstrates a mildly quadratic relationship between sequence length and computational cost, that does not represent a real burden even a the longest sequences considered.

These computational achievements were supported by robust experimental validation. Two assays were conducted: the first assessed the feasibility of CertPrime designs for eight sequences, all of which yielded the expected results without spurious interactions. The second assay compared designs of the S100A4 gene produced by CertPrime and other tools. CertPrime consistently outperformed its competitors, delivering a correct, dimer-free design, whereas all other tools failed to meet these criteria. Even if there is no guarantee that CertPrime finds the best global design, nor that an arbitrary sequence can be synthesised effectively, our results support the value of the tool in typical cases, and its superiority over other available tools. In conclusion, CertPrime represents a significant advancement in oligonucleotide design for gene synthesis, combining computational efficiency, experimental reliability, and unparalleled flexibility to address the challenges of modern genomic applications.

## Acknowledgments

The authors express their gratitude to all the members of Certest Biotec S.L.

## Funding

Research reported in this publication was supported by Gobierno de Aragón (Spain) through project IDMF/2021/0009 (Nuevas tecnoloǵıas para el diseño y obtención de vacunas de ARN en Araǵon) and E30 20R (“Su- percomputación Y Física De Sistemas Complejos Y Bioĺogicos” – COM- PHYS). P.B. acknowledges support through grant PID2020-113582GB-I00 and PID2023-147734NB-I00, funded by MCIN/AEI/10.13039/ 501100011033.

## Declaration of competing interests

Juan Martínez-Oliván, Esther Broset, Fadi Hamdan, Irene Blasco-Machín, David Luna-Cerralbo and Ana Serrano are employees at Certest Biotec S.L.

## Code and data availability

The sequences for the proteins reported in Table 3 and their %CG profiles are included in the Supplementary Information. The oligo designs used in Figure 8 are included as Supplementary Files. The CertPrime code is available from the corresponding authors PB and EB on reasonable request.

## CRediT authorship contribution statement

FH, JMO and PB conceived, designed and supervised the study. PB and DLC conceived, implemented and interpreted the computational approach. EB, IBM and AS conceived and interpreted the experimental methods. AS performed synthesis and PCR experiments. DLC wrote the initial draft manuscript. EB and PB wrote the final version manuscript. JMO and PB provided the funding. All authors contributed to the article and approved the submitted version.

## Declaration of Generative AI and AI-assisted technologies in the writing process

During the revision of this work, the authors used DeepL in order to improve the language quality of some sentences. After using this tool, the authors reviewed and edited the content as needed, rejecting the suggestions that did not reflect the original meaning of the sentences. The authors take full responsibility for the content of the publication.

## Appendix A: Supplementary information

Supplementary results can be found online at the the corresponding link.

