## Supplementary figures and data for "CertPrime: a new oligonucleotide design tool for gene synthesis"

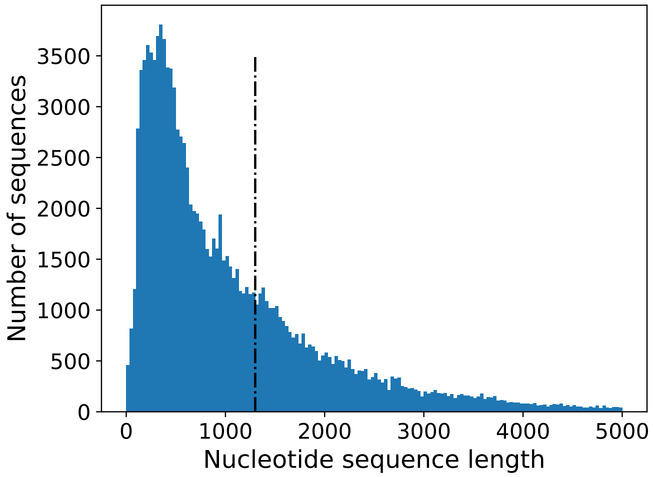


Figure 1: **Distribution of nucleotide sequence lengths in the human genome downloaded from Ensembl**. The bulk of the sequences are concentrated in lengths from 200 to 2000 nts, indicating a substantial pool with characteristics similar to those of the sequences typically addressed in biotechnological applications. The vertical line marks the maximum length of sequence we took for the DS1 test.

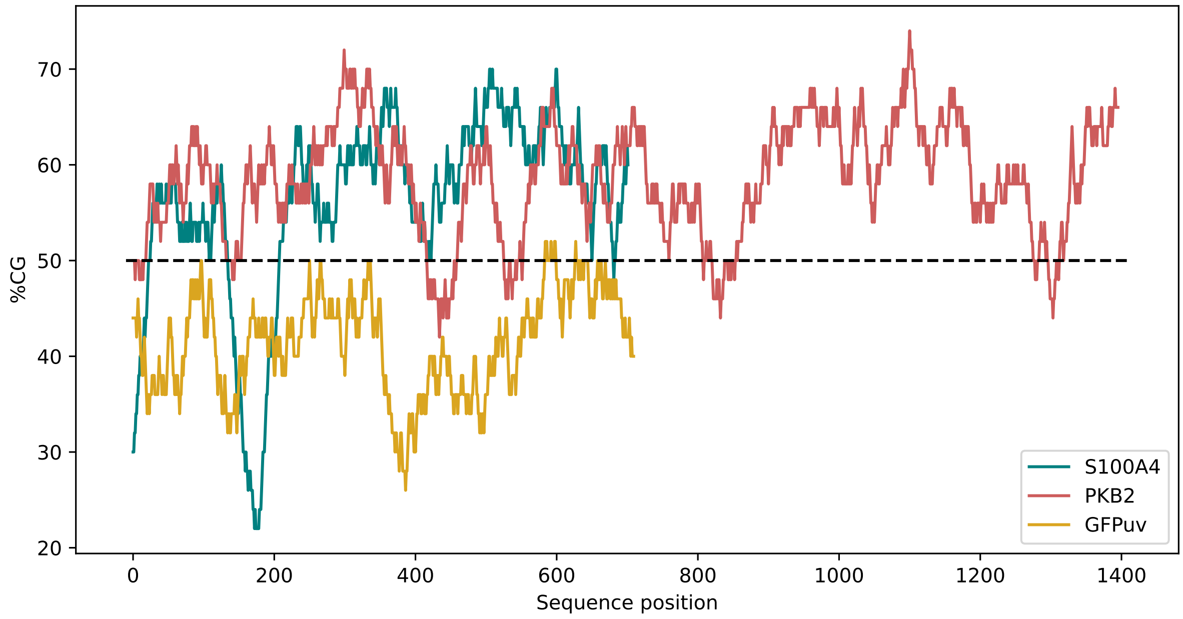


Figure 2: **GC content profiles of three nucleotide sequences, PKB2 (1446 nts), GFPuv (760 nts), and S100A4 (752 nts), calculated using 50-nucleotide sliding windows.** The window size of 50 nts was chosen because it is sufficiently large that excessively high or low GC content can present a challenge when designing oligonucleotides.

**Nucleotide sequences of the S100A4, PKB2, and GFPuv genes:**

```
>PKB2 gene
ATGAATGAGGTGTCTGTCATCAAAGAAGGCTGGCTCCACAAGCGTGGTGAATACATCAAGACCTGGAGGCCACGGTACTTCCTGCTGAAGAGCGA
CGGCTCCTTCATTGGGTACAAGGAGAGGCCCGAGGCCCCTGATCAGACTCTACCCCCCTTAAACAACTTCTCCGTAGCAGAATGCCAGCTGATGA
AGACCGAGAGGCCGCGACCCAACACCTTTGTCATACGCTGCCTGCAGTGGACCACAGTCATCGAGAGGACCTTCCACGTGGATTCTCCAGACGAG
AGGGAGGAGTGGATGCGGGCCATCCAGATGGTCGCCAACAGCCTCAAGCAGCGGGCCCCAGGCGAGGACCCCATGGACTACAAGTGTGGCTCCCC
CAGTGACTCCTCCACGACTGAGGAGATGGAAGTGGCGGTCAGCAAGGCACGGGCTAAAGTGACCATGAATGACTTCGACTATCTCAAACTCCTTG
GCAAGGGAACCTTTGGCAAAGTCATCCTGGTGCGGGAGAAGGCCACTGGCCGCTACTACGCCATGAAGATCCTGCGAAAGGAAGTCATCATTGCC
AAGGATGAAGTCGCTCACACAGTCACCGAGAGCCGGGTCCTCCAGAACACCAGGCACCCGTTCCTCACTGCGCTGAAGTATGCCTTCCAGACCCA
CGACCGCCTGTGCTTTGTGATGGAGTATGCCAACGGGGGTGAGCTGTTCTTCCACCTGTCCCGGGAGCGTGTCTTCACAGAGGAGCGGGCCCGGT
TTTATGGTGCAGAGATTGTCTCGGCTCTTGAGTACTTGCACTCGCGGGACGTGGTATACCGCGACATCAAGCTGGAAAACCTCATGCTGGACAAA
GATGGCCACATCAAGATCACTGACTTTGGCCTCTGCAAAGAGGGCATCAGTGACGGGGCCACCATGAAAACCTTCTGTGGGACCCCGGAGTACCT
GGCGCCTGAGGTGCTGGAGGACAATGACTATGGCCGGGCCGTGGACTGGTGGGGGCTGGGTGTGGTCATGTACGAGATGATGTGCGGCCGCCTGC
CCTTCTACAACCAGGACCACGAGCGCCTCTTCGAGCTCATCCTCATGGAAGAGATCCGCTTCCCGCGCACGCTCAGCCCCGAGGCCAAGTCCCTG
CTTGCTGGGCTGCTTAAGAAGGACCCCAAGCAGAGGCTTGGTGGGGGGCCCAGCGATGCCAAGGAGGTCATGGAGCACAGGTTCTTCCTCAGCAT
CAACTGGCAGGACGTGGTCCAGAAGAAGCTCCTGCCACCCTTCAAACCTCAGGTCACGTCCGAGGTCGACACAAGGTACTTCGATGATGAATTTA
CCGCCCAGTCCATCACAATCACACCCCCTGACCGCTATGACAGCCTGGGCTTACTGGAGCTGGACCAGCGGACCCACTTCCCCCAGTTCTCCTAC
TCGGCCAGCATCCGCGAGTGA
>GFPuv gene
AGAGGATCCCCGGGTACCGGTAGAAAAAATGAGTAAAGGAGAAGAACTTTTCACTGGAGTTGTCCCAATTCTTGTTGAATTAGATGGTGATGTTA
ACGGGCACAAATTTTCTGTCAGTGGAGAGGGTGAAGGTGATGCAACATACGGAAAACTTACCCTTAAATTTATTTGCACTACTGGAAAACTACCT
GTTCCATGGCCAACACTTGTCACTACTTTCTCTTATGGTGTTCAATGCTTTTCCCGTTATCCGGATCATATGAAACGGCATGACTTTTTCAAGAG
TGCCATGCCCGAAGGTTATGTACAGGAACGCACTATATCTTTCAAAGATGACGGGAACTACAAGACGCGTGCTGAAGTCAAGTTTGAAGGTGATA
CCCTTGTTAATCGTATCGAGTTAAAAGGTATTGATTTTAAAGAAGATGGAAACATTCTCGGACACAAACTCGAGTACAACTATAACTCACACAAT
GTATACATCACGGCAGACAAACAAAAGAATGGAATCAAAGCTAACTTCAAAATTCGCCACAACATTGAAGATGGAAGCGTTCAACTAGCAGACCA
TTATCAACAAAATACTCCAATTGGCGATGGCCCTGTCCTTTTACCAGACAACCATTACCTGTCGACACAATCTGCCCTTTCGAAAGATCCCAACG
AAAAGCGTGACCACATGGTCCTTCTTGAGTTTGTAACTGCTGCTGGGATTACACATGGCATGGATGAGCTCTACAAATAATGAATTCCAACTGAG
>S100A4 gene
GTTTTTGTTTCTGAATCTTTATTTTTTTAAGAGACAAGGTCCTCTGTGTTGCTCAGGCTGGAGAGCAGTGGCTTGAGCATAGCCAACTGCAGTCT
CGAACTCCTGGGCTCAAATGATCCTCCTGTCTCAGCTTCCTGACTAGCTGGGACTACAGGCTACAGCCATGCTGCCCAGCTAATTAAAAAAAAAA
```

ATTGTTTTTCCTTTTTATAGAGACAGAAGTCTCTCTATGTTGCCTAGGCTGGTCTTGAACTCCTGGCCTCAGGCGATCCTCCCATCTCCCCCCTA
GCTTTTGTGTCACCACATTTCCAGGGCAATCTCCCACCTGTCACCCACCACCCCCTGCATCTCCTTTCCTAGGTCCCCATGGGACTACTCCCTGT
CCCCCATGCTCCAGGCACAGGCTGCCCCTTCCTCCACCTCTCTAAAACTCAGGCTGAGCTATGTACACTGGGTGGTGCCCATCTCATCCAGTCCC
CTGCTAGTAACCGCTAGGGCTTACCCGTTACCCACGGGTGCCCACCTGGGAACAGGAGGCTTGGTTCCACGGCTGGGCTGGTGGAGGGTGCTGTG
GCACTTACCGCATCAGCCCACAGCAGGAAGGCAGTATCCGCTCTCCCCTGTCCCCTGCTATGGGCAGGGCCTGGCTGGGGTATAAATAGGTCAGA
CCTCTGGGCCGTCCCCATTCTTCCCCTCTCTACAACCCTCTCTCCTCAGCGCTTCTTCATCAAGATCTGGCCTCGGCGGCCAAGCTT
